# Comparative genomics of *Streptococcus oralis* identifies large scale homologous recombination and a genetic variant associated with infection

**DOI:** 10.1101/2022.08.05.502949

**Authors:** Luke R. Joyce, Madison A. Youngblom, Harshini Cormaty, Evelyn Gartstein, Katie E. Barber, Ronda L. Akins, Caitlin S. Pepperell, Kelli L. Palmer

## Abstract

The viridans group streptococci (VGS) are a large consortium of commensal streptococci that colonize the human body. Many species within this group are opportunistic pathogens causing bacteremia and infective endocarditis (IE), yet little is known about why some strains cause invasive disease. Identification of virulence determinants is complicated by the difficulty of distinguishing between the closely related species of this group. Here, we analyzed genomic data from VGS isolated from patient blood cultures with invasive infections and from oral swabs of healthy volunteers and determined the best performing methods for species identification. Using whole-genome sequence data, we characterized the population structure of a diverse sample of *Streptococcus oralis* isolates and found evidence of frequent recombination. We used multiple genome-wide association study tools to identify candidate determinants of invasiveness. These tools gave consistent results, leading to the discovery of a single synonymous single nucleotide polymorphism (SNP) that was significantly associated with invasiveness. This SNP is within a previously undescribed gene that is conserved across the majority of VGS species. Using growth in the presence of human serum and a simulated infective endocarditis vegetation model, we were unable to identify a phenotype for the enriched allele in laboratory assays, suggesting a phenotype may be specific to natural infection. These data highlight the power of analyzing natural populations for gaining insight into pathogenicity, particularly for organisms with complex population structures like the VGS.

**Importance:** The viridians group streptococci (VGS) are a large collection of closely related commensal streptococci, with many being opportunistic pathogens causing invasive diseases such as bacteremia and infective endocarditis. Little is known about virulence determinants in these species, and there is a distinct lack of genomic information available for the VGS. In this study, we collected VGS isolates from invasive infections and healthy volunteers and performed whole genome sequencing for a suite of downstream analyses. We focused on a diverse sample of *Streptococcus oralis* genomes and identified high rates of recombination in the population as well as a single genome variant highly enriched in invasive isolates. The variant lies within a previously uncharacterized gene, *nrdM*, which shares homology with the anaerobic ribonucleoside triphosphate reductase, *nrdD*, and is highly conserved among VGS. This work increases our knowledge of VGS genomics and indicates that differences in virulence potential among *S. oralis* isolates is, at least in part, genetically determined.

## Introduction

The viridans group streptococci (VGS) comprise a diverse collection of alpha and non-hemolytic streptococci that inhabit the oral cavity and gastrointestinal and genitourinary tracts of healthy humans (*1*). VGS are also associated with invasive disease, particularly in immunocompromised hosts, and are estimated to cause ~23% of Gram-positive bacteremia (*2*, *3*) and ~17% of infective endocarditis (IE) cases (*4*). Bacterial determinants of invasiveness among the VGS are not well understood. Research in this area is hampered by the fact that the specific species of VGS causing bacteremia and IE are infrequently determined in a clinical context due to lack of resolution of existing diagnostic microbiological tools (*5*, *6*). VITEK®2 and MicroScan allow for general assignment of isolates to VGS, and assignment of a limited number of VGS species specifically. Even a relatively newer technique in clinical diagnostics, matrix assisted laser desorption ionization-time of flight mass spectrometry (MALDI-TOF MS), fails to resolve certain VGS species, including *Streptococcus mitis* and *Streptococcus oralis* (*7*). 16S rRNA gene sequencing, multilocus sequence analysis and other genotyping schemes (*5–10*), and GyrB typing (*11*) are commonly used methods for molecular VGS identification but are not generally employed in clinical laboratories.

Retrospective studies using molecular approaches have determined that in addition to being present as oral commensals in healthy individuals, *S. mitis* and *S. oralis* stand out among VGS as major causative agents of VGS bacteremia and IE (*8*, *12–15*). *S. mitis* and *S. oralis* are members of the mitis group streptococci, a subgroup within the VGS that is closely related to the major human pathogen *Streptococcus pneumoniae* (*16, 17*). A recent study in oncology patients demonstrated that *S. mitis* (58%) and *S. oralis* (19%) were the most frequently identified species in VGS infections over ~1.5 years (*8*). While *S. mitis* and *S. oralis* are a significant burden on immunocompromised patients, the mechanisms of virulence within these species have not been fully elucidated. More specifically, it is not known whether all members of these species have equal pathogenic potential, or whether some strains have a higher propensity for causing invasive disease than others.

In this study, we aimed to investigate the mechanisms of invasiveness among VGS by characterizing species diversity of presumptive VGS obtained clinically from bacteremia and endocarditis patients. Clinical isolate genomes obtained in this study were supplemented by existing genome sequences with curated metadata in public databases and genome sequences of oral isolates collected from healthy volunteers. Our results support metagenomic sequence binning as a high-resolution tool for differentiating species within the VGS grouping. After a preliminary analysis in which we determined species designations for clinical isolates that were diagnosed as VGS, we focused our study on *S. oralis* as a prominent cause of invasive infection. Within a large sample of *S. oralis* isolates from all three described sub-species (*18*) (*subsp. dentisani, subsp. tigurinus*, and *subsp. oralis*) we found high levels of diversity and strikingly high recombination rates. We used multiple genome-wide association study (GWAS) methods to test the hypothesis that specific genetic variations are associated with invasive infection (when compared to commensal isolates) among *S. oralis* isolates. We discovered a SNP in a previously uncharacterized gene that is significantly enriched in invasive isolates compared to non-invasive isolates. The contribution of this novel locus to growth with human serum and in a simulated infective endocarditis vegetation model (*19*) was assessed, although we were unable to identify a phenotype for a gene knockout or for either allele of the significant variant under the conditions tested. This work has 1) increased the genomic information available for VGS strains, 2) described population structure and large-scale homologous recombination within the *S. oralis* species, and 3) provided evidence that the propensity for virulence in *S. oralis* is at least in part genetically determined.

## Results

### GyrB typing and Kraken efficiently speciate *S. mitis* and *S. oralis*

VGS isolates from Dallas, Texas and Jackson, Mississippi areas were collected from clinically confirmed VGS bacteremia and infective endocarditis patients. Isolates were initially characterized using either VITEK®2 or MicroScan platforms. A total of 66 clinical isolates were successfully sub-cultured in the laboratory, however two isolates were incorrectly identified as *Streptococcus* spp.: one *Enterococcus faecalis*, and one *Aerococcus urinae* strain, resulting in 64 presumptive VGS strains collected (Table S1). To compare isolates causing invasive disease against commensal isolates, healthy volunteers were recruited for oral swab collection on the University of Texas at Dallas campus, for a total of 81 VGS isolates (Table S1).

As has previously been reported, clinical methods of VGS species identification (VITEK®2 and MicroScan) were not effective for delineating closely related species, particularly within the mitis group; however, they were almost always correct in identifying an isolate as a part of the VGS (Table S1). The taxonomy of the clinical and oral isolates was analyzed by 16S rRNA gene sequencing, GyrB typing (*11*), and analysis of Illumina whole genome sequencing (WGS) data using Kraken (*20*). While we expected that WGS would provide the clearest results, we also wanted to assess the utility of the GyrB typing scheme for a diverse set of VGS species, as this method is cheaper, faster and more feasible for researchers without computational experience and resources. Using these three different methods of species identification, we classified isolates into VGS groups using the taxonomy described by Facklam (*21*). All six of the major VGS groups (*1*) were represented in our sample of 81 isolates, with two isolates not fitting into any of these six groups (Figure 1A). 16S rRNA sequencing was frequently unable to provide resolution to the species level but was often able to identify which group the isolate belonged to. GyrB typing (*11*) and Kraken (*20*) were both effective in distinguishing *S. oralis* and *S. mitis*, however GyrB typing is not generally accurate for other VGS groups (Figure 1A).

**Figure 1:**
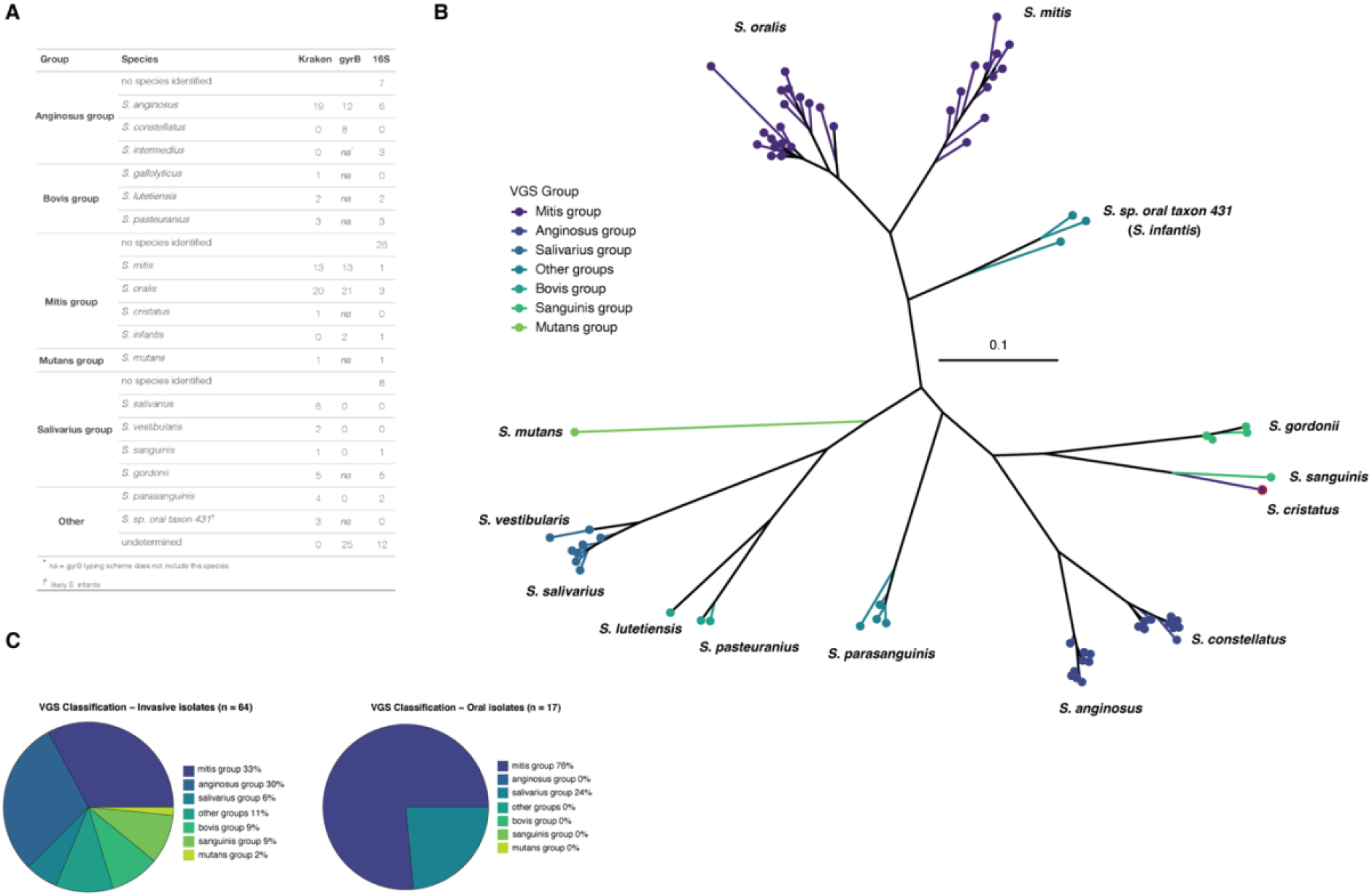
Species identification of blood and oral commensal viridans group streptococci (VGS). **A)** Three different methods for species identification were used (Kraken, GyrB typing & 16S rRNA sequencing) providing differing results. “na” values in the gyrB column indicate species not included in this typing scheme. “No species identified” indicates that the isolate could only be identified to the group level. **B)** Phylogenetic tree of all 81 isolates inferred from an alignment of GyrB sequences and identified by VGS group and species as assigned by Kraken. The isolate which appears to have been mis-identified by Kraken is outlined in red. **C)** Invasive (left) and oral (right) isolates by VGS group according to species identification performed by Kraken.

One downside of species identification with Kraken is it seems unable to distinguish between anginosus group isolates: *S. anginosus* and *S. constellatus* are species within the anginosus group (also known as the milleri group) of the VGS and are recognized as abscess-causing bacteria, with more recent data suggesting their emergence as uropathogens (*1*, *22*, *23*). Kraken identified 19 *S. anginosus* isolates, and via GyrB typing a total of 20 anginosus group isolates were identified (12 *S. anginosus* and 8 *S. constellatus*). These data suggest that *S. mitis* and *S. oralis* are accurately speciated with either Kraken or GyrB typing, yet anginosus group species may be better identified by GyrB typing.

One drawback to species identification with GyrB typing is the current scheme is limited in the species it can distinguish (Figure 1A). To further assess the relative functionality of these two methods we inferred a phylogenetic tree from GyrB sequences and mapped species as defined by Kraken onto the tree (Figure 1B). The added benefit of a phylogeny of GyrB sequences is the visualization of additional species not included in the typing scheme (e.g. species in the bovis group; Figure 1B) which when combined with Kraken output allowed us to confidently assign species to most isolates. There was only a single instance where Kraken may have mis-identified the species: an isolate identified by Kraken as *S. cristatus* (a member of the mitis group) appears to cluster with *S. sanguinis* isolates (of the sanguinis group) on the GyrB phylogeny (Figure 1B). Additionally, combining data from Kraken and GyrB typing showed that 2 out of the 3 isolates identified as *S. sp. oral taxon 431* by Kraken were identified as *S. infantis* (a member of the mitis group) by GyrB typing, and all 3 cluster together on the GyrB phylogeny, indicating that *S. infantis* is likely the correct designation (Figure 1B).

Overall, our data show that closely related *S. mitis* and *S. oralis* can be speciated accurately via either Kraken or GyrB typing; however, anginosus group isolates are better distinguished by GyrB typing. For datasets suspected to contain a diverse sample of different VGS species, species identification using Kraken with whole genome sequence data appears the most robust. GyrB typing is an accurate method for making distinctions within species subsets, and a phylogeny of GyrB sequences may also provide additional information. Our results support previous assertions that 16S rRNA sequencing is not an effective method for distinguishing the closely related species of the VGS (*1*).

When we separated our sample into isolates from blood cultures and those from the mouths of healthy persons, we saw that invasive isolates spanned all 6 groups while isolates from oral isolates were less diverse and only originated from the mitis and salivarius groups (Figure 1C). The lack of diversity in the oral isolates is due to the use of Mitis-Salivarius Agar which inhibits Gram-negative and most Gram-positive bacterial growth due to the presence of inhibitory nutrients; however, *S. mitis, S. salivarius*, and enterococci will still grow and produce different colony morphologies to allow preferential selection (*24*). It has been reported that *S. mitis* is the predominant species found in healthy oral microbiomes (*25–30*) with other commonly found VGS species including *S. oralis* (*27, 28*), *S. sanguinis* (*28, 29*), and *S. salivarius* (*29*).

### High diversity among commensal and invasive *S. oralis* strains

*S. oralis* was the predominant single species (20/81 isolates) in our sample, and so to identify possible genetic variants associated with invasive infection, we gathered a larger sample of *S. oralis*. We obtained all isolates labeled as *S. oralis* and *S. mitis* from NCBI (see Methods for inclusion criteria) and used Kraken, a core genome phylogeny, and accessory gene content to confirm *S. oralis* isolates and identify mislabeled isolates. We found that a core genome alignment produced by Roary (*31*) effectively delineates *S. oralis* from *S. mitis*, as do patterns of accessory gene content (Figure S1). We were able to identify 11 isolates mis-labeled as *S. mitis* within NCBI databases, which we identified as *S. oralis* using these methods (Table S2). We ended up with a total of 108 *S. oralis* isolates: 57 oral commensals and 51 from invasive infections (including bacteremia and IE) that we refer to as “oral” and “blood” isolates, respectively (Table S2). After assembly and annotation of all genomes, we performed a pangenome analysis with Roary and found that our sample has a core genome of only 801 genes – with an average genome size of 1898 genes. This means that a significant proportion of genes (57%) encoded by an individual isolate are variable accessory genes (Figure 2A). The accessory gene content in our sample of *S. oralis* isolates is highly diverse, with 88% of all genes found in our sample at a frequency of 14% or less, and > 19,000 genes identified in the pan genome (Figure 2A). This high level of gene content diversity is mirrored in the core genome phylogeny, which has long terminal branch lengths indicating high variability between core genome sequences (Figure 2B). Isolates from commensal and invasive sources are interspersed on the phylogeny and do not generally form monophyletic clades, nor are the commensal or invasive phenotypes associated any sub-clades within the phylogeny (Figure 2B).

**Figure 2:**
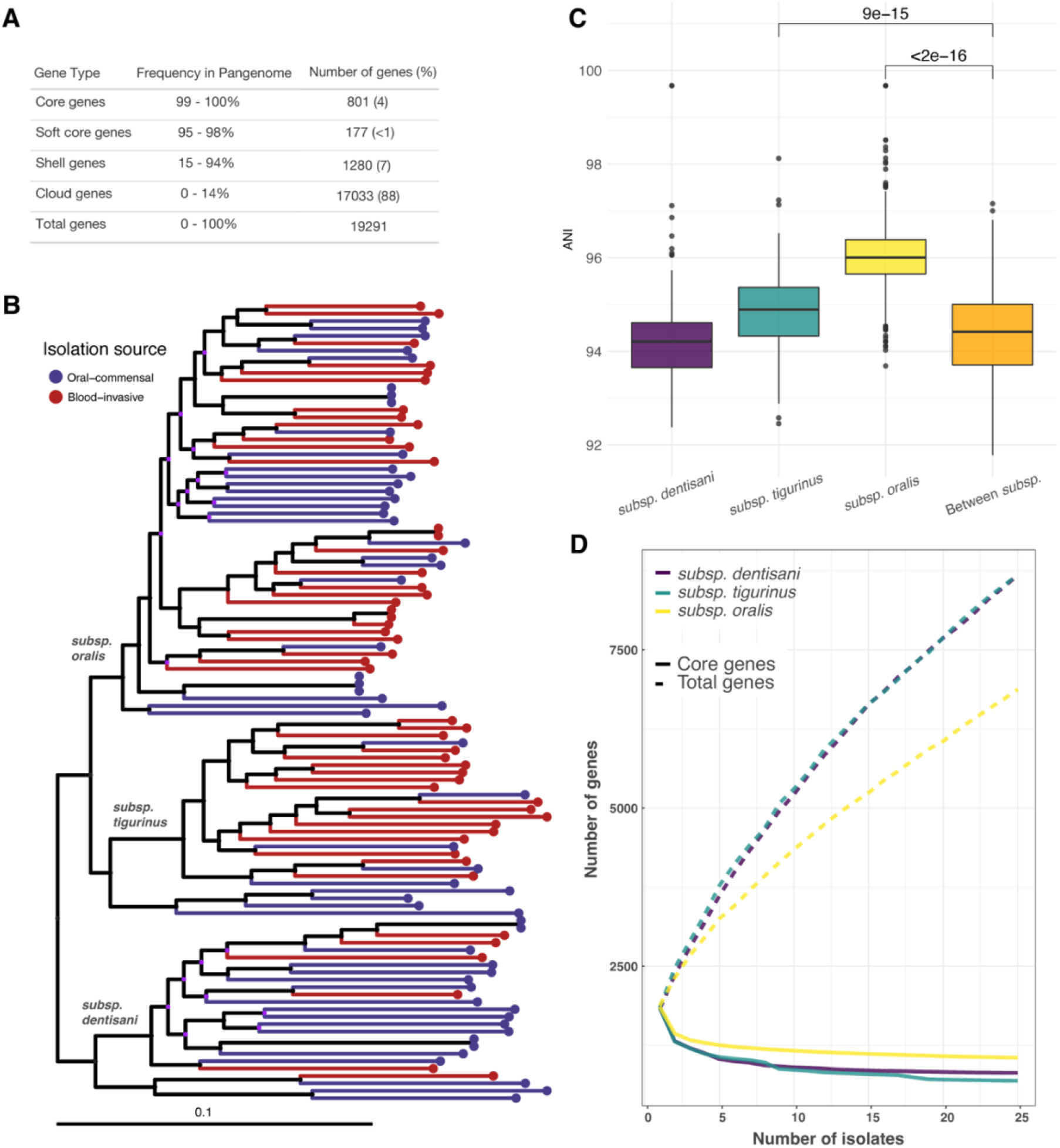
Pangenome analysis of *S. oralis* strains. **A)** Summary statistics from pangenome analysis of all *S. oralis* GWAS isolates (n=108). **B)** Core genome phylogeny of *S. oralis* isolates inferred using RAxML and midpoint rooted. Tips colored by isolation source (oral-commensal in blue, blood-invasive in red). 108 isolates total, 57 isolated from the oral cavity and 51 isolated from invasive infections. Three subspecies are labeled and scale bar given in SNPs per site. Nodes colored in purple represent bootstrap values < 50. **C)** Average nucleotide identity (ANI) calculated within and between the three subspecies of *S. oralis* phylogeny show relatively low sequence conservation (~94-96% ANI) between even strains of the same subspecies. ANI values within *subsp. oralis* and *subsp. tigurinus* show significantly higher sequence conservation within compared to between subspecies (Mann-Whitney U test with Benjamini-Hochberg correction). **D)** Accumulation and rarefaction curves for three subspecies. All samples were repeatedly sub-sampled to the size of the smallest sample (*subsp. dentisani*, n=25) and the median core and total gene values plotted for 100 iterations.

We identified three sub-clades within our core genome phylogeny (Figure 2B) that we found to correspond with the three previously described sub-species of *S. oralis: subsp. dentisani, subsp. tigurinus*, and *subsp. oralis* (*18*) by cross-referencing our tree with the subspecies of some of the publicly available isolates in our dataset (Table S2). Average nucleotide identity (ANI) values for core genome sequences within each subspecies show that *subsp. oralis* is the most conserved, followed by *subsp. tigurinus* and *subsp. dentisani* (Figure 2C). *Subsp. dentisani* is unique in that it has ANI values resembling those measured between isolates of different subspecies – this indicates that levels of diversity within this subspecies are similar to those found between subspecies (Figure 2C). Following the trends we identified in core genome diversity levels, the pangenome of *subsp. oralis* isolates appears more conserved (more core genes, fewer accessory genes) than either *subsp. tigurinus* or *subsp. dentisani* (Figure 2D). Higher core genome and pangenome diversity levels among *subsp. tigurinus* and *subsp. dentisani* isolates could indicate that these isolates have access to more diverse partners for horizontal gene transfer (HGT).

### High levels of recombination among *S. oralis* isolates

Viridans group streptococci (VGS) are known for being naturally competent (*32*) and for evolving rapidly via widespread homologous recombination (*33*, *34*). We characterized the signatures of recombination in the core genome of *S. oralis* using Gubbins (*35*) and identified extreme amounts of recombination: 99.9% of the core genome is within a predicted recombinant fragment in at least one isolate (Figure 3A). Additionally, we noted that recombinant fragments are not shared across multiple isolates but are usually present in only a handful of isolates (Figure 3B). Using ClonalFrameML (*36*) we calculated the *r/m* value - the ratio of SNPs imported via recombination (r) to those introduced randomly (m) – and found that with an *r/m* of 5.77, genetic diversity in our sample is ~6x more likely to be introduced via recombination. This is slightly less than the notoriously recombinogenic *S. pneumoniae* [*r/m* = ~7; (*33*)] but much higher than other IE causing bacteria such as *S. aureus* [*r/m* = <1; (*37*)].

**Figure 3:**
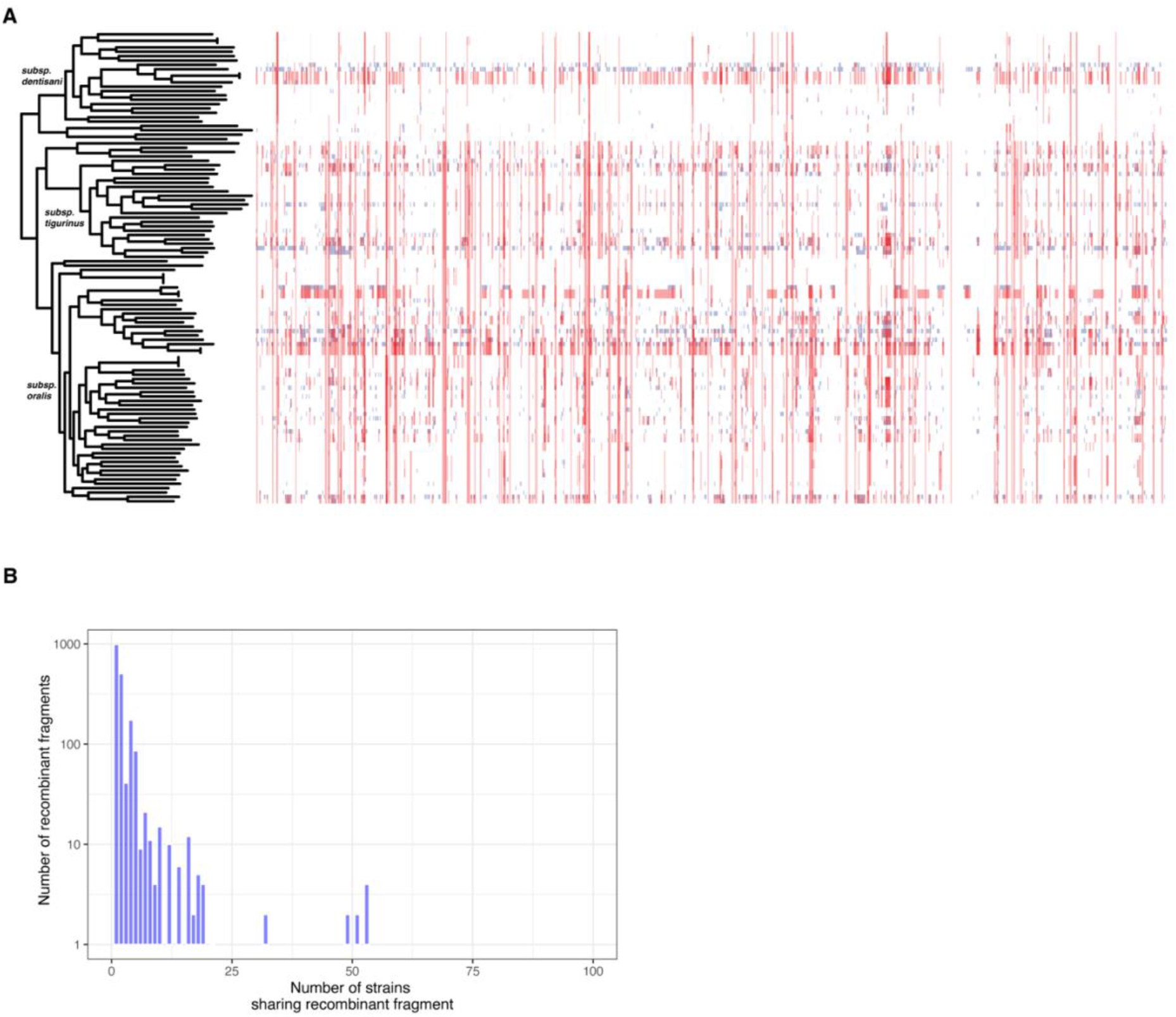
*S. oralis* strains are exceptionally recombinogenic. **A)** Recombination tracts predicted by Gubbins in the core genome of *S. oralis* isolates plotted alongside the core genome phylogeny. Red tracts represent recombination within the sample and blue tracts represent recombination with isolates outside this sample. 99.9% of the core genome has been affected by recombination. Visualization performed with Phandango. **B)** Histogram showing the number of strains in our sample that share a given recombinant fragment. The x-axis is the number of strains that share a recombinant fragment (with a maximum of 108 isolates, the size of our sample) and the y-axis is the number of fragments identified by Gubbins that are shared by that number of isolates.

### Genome wide association study reveals variant associated with invasiveness

With a large sample balanced between our phenotypes of interest, we used multiple genome-wide association study (GWAS) methods to identify genetic variants associated with invasive disease, which included isolates from both IE and bacteremia patients. We started by looking for associations between accessory genes and invasiveness using Scoary (*38*), which yielded no significant results. This is perhaps not surprising given that accessory gene content is so diverse in the sample and individual accessory genes are thus unlikely to be shared by a large proportion of isolates (Figure 1A).

To identify core genome variants associated with invasiveness, we started with a F_ST_ outlier analysis, which delineates allele frequency differences between sub-populations and identifies variants with extreme measures of differentiation. We defined sub-populations of our sample as being of “oral” or “blood” source and used vcflib (https://github.com/vcflib/vcflib) to calculate Weir and Cockerham’s F_ST_ (wcF_ST_, abbreviated to F_ST_) for bi-allelic SNPs (n=67,026) in the core genome alignment (Figure 4A). We identified a single SNP with a significant F_ST_ value, indicating significant allele frequency differences between oral and blood sub-populations of *S. oralis* (Figure 4A). The SNP of interest lies within a gene originally annotated as a homologue of *nrdD*, an anaerobic ribonucleoside triphosphate reductase present in many streptococci, involved in synthesis of deoxyribonucleotides under anaerobic conditions (*39*). However, further inspection of the sequence revealed that the canonical *nrdD* gene was annotated separately in our *S. oralis* isolates, and that the novel protein was shorter than *nrdD*. The novel protein does share some sequence features with *nrdD* including an ATP cone domain, and so we will refer to the novel locus as *nrdM*. Given the amount of recombination present in our sample (Figure 3A) we validated the results of our F_ST_ outlier analysis using GWAS methods specifically designed for use with microbial genomes. We used two programs: treeWAS (*40*), which corrects for the presence of recombination, and BugWAS (*41*), which identifies lineage effects and controls for population structure. Results from both of these tools replicated the results of our F_ST_ outlier analysis with both methods returning a single variant that was significantly associated with the phenotype: the same SNP in *nrdM* (Figure S2).

**Figure 4:**
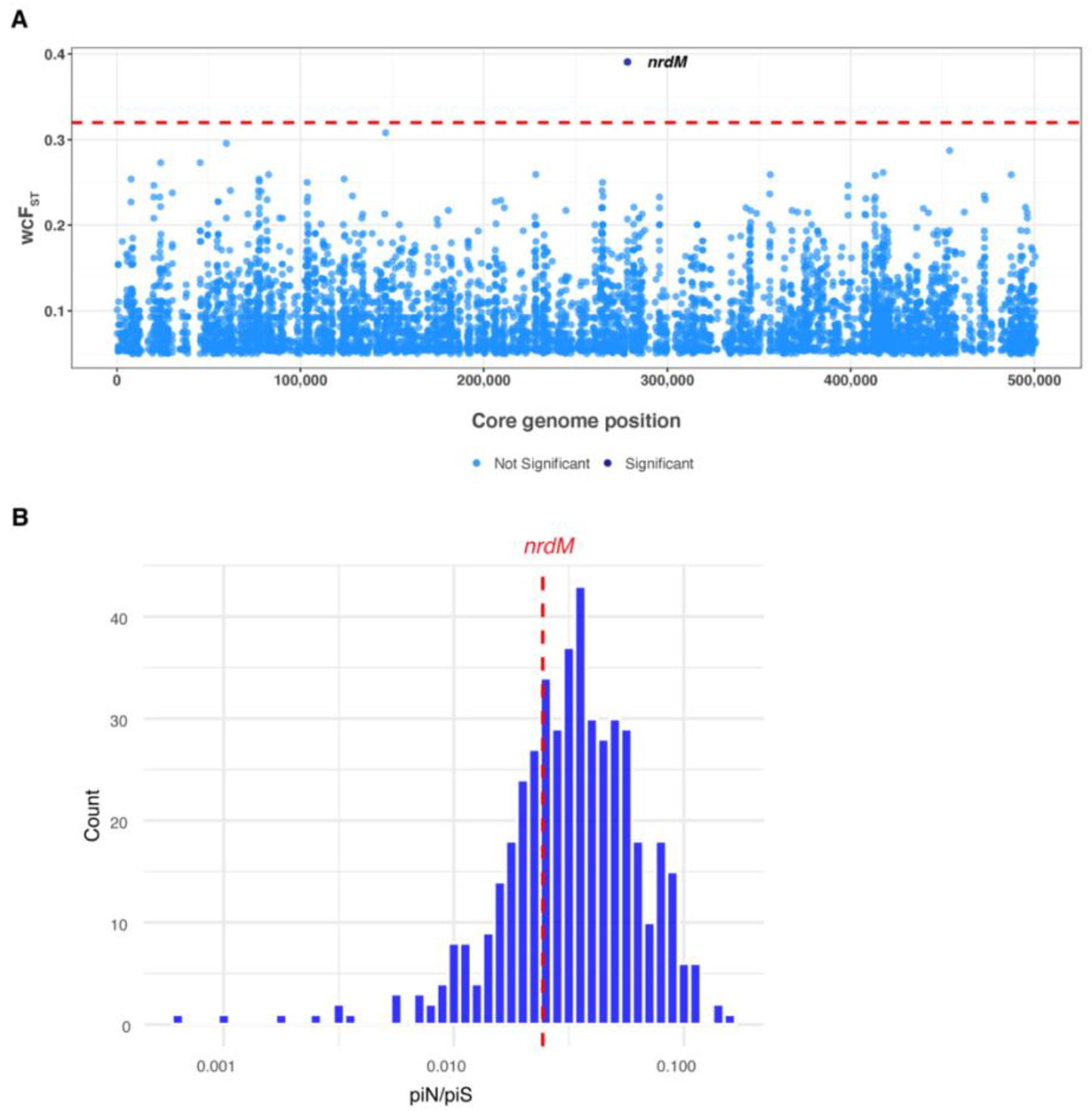
A Single core genome variant is associated with invasiveness in *S. oralis* isolates. **A)** Weir and Cockerham’s F_ST_ (wcF_ST_) values calculated for each core genome variant (n=67,026) and plotted against the core genome position. Significance threshold (red dotted line) estimated by taking the maximum wcFst value from 100 random permutations of phenotypes. Variants with non-significant F_ST_ values shown in light blue, a single variant with a significant F_ST_ value shown in dark blue. **B)** Pairwise piN/piS values calculated and averaged for each core gene and plotted as a histogram. Average piN/piS value across all genes is 0.039, red dotted line represents the average piN/piS value for *nrdM* (0.024).

The SNP of interest in *nrdM* is a synonymous mutation (I78I), and mapping the nucleotide alleles of the mutation (C233T) onto the core genome phylogeny shows interspersal of alleles on the tree, with some structure in *subsp. dentisani* and *subsp. tigurinus* (Figure S3). Most interesting is the striking association between isolates from invasive infections and the *nrdM* allele: out of 51 invasive isolates in our sample, 39 encode the *nrdM* SNP associated with invasiveness (Figure S3; Table S2). A multiple sequence alignment of NrdM from our sample of *S. oralis* strains reveals high protein sequence conservation (Figure S4), and so to determine whether *nrdM* is more or less conserved than other core genes we calculated the average πN/πS (piN/piS) values for each core gene (n=801). The distribution of gene-wise values of piN/piS indicate the majority of genes are evolving under relatively strong purifying selection, including *nrdM*, which had a piN/piS value slightly below the mean (Figure 4B). This is not surprising given that recombination strengthens the efficiency of selection by enabling rapid removal of deleterious mutations (*42*).

### Selection on *nrdM* variant

Given the strong association between the *nrdM* variant and invasiveness, we wondered if this SNP is under positive selection in our sample. One method for identifying positive selection is the identification of homoplastic mutations, *i.e*. mutations that arose more than once on the phylogeny, which we have used previously to screen for drug resistance loci in *Mycobacterium tuberculosis* (*43*). We used TreeTime (*44*) to identify homoplastic mutations in our sample and found numerous homoplastic mutations – this is expected since intergenomic recombination can produce homoplasies by lateral transfer of sequence variants. The variant in *nrdM* arose 20 times on the phylogeny, which was in the 85^th^ percentile of mutation multiplicity (Figure S5). This finding, along with gene-wise piN/piS values (Figure 4B), suggests that selection pressures are similar at this locus to others in the genome; however, further analysis with a larger sample size could elucidate more subtle signs of selection.

### *nrdM* is conserved among VGS species

In order to investigate the presence of *nrdM* homologues in other *Streptococcus* spp. we used Blast to search annotated genes in all newly sequenced isolates and found *nrdM* to be present in all VGS species in our original sample, except for two bovis group species: *S. lutetiensis* and *S. pasteuranius* (Figure 5A). Additionally, the position of the variant associated with invasiveness in *S. oralis* is conserved, and we were able to identify variation in *nrdM* alleles between different species. As in *S. oralis*, NrdM is highly conserved at the protein level across the VGS species in our sample, indicating the gene may have a conserved function across multiple species (Figure 5B).

**Figure 5:**
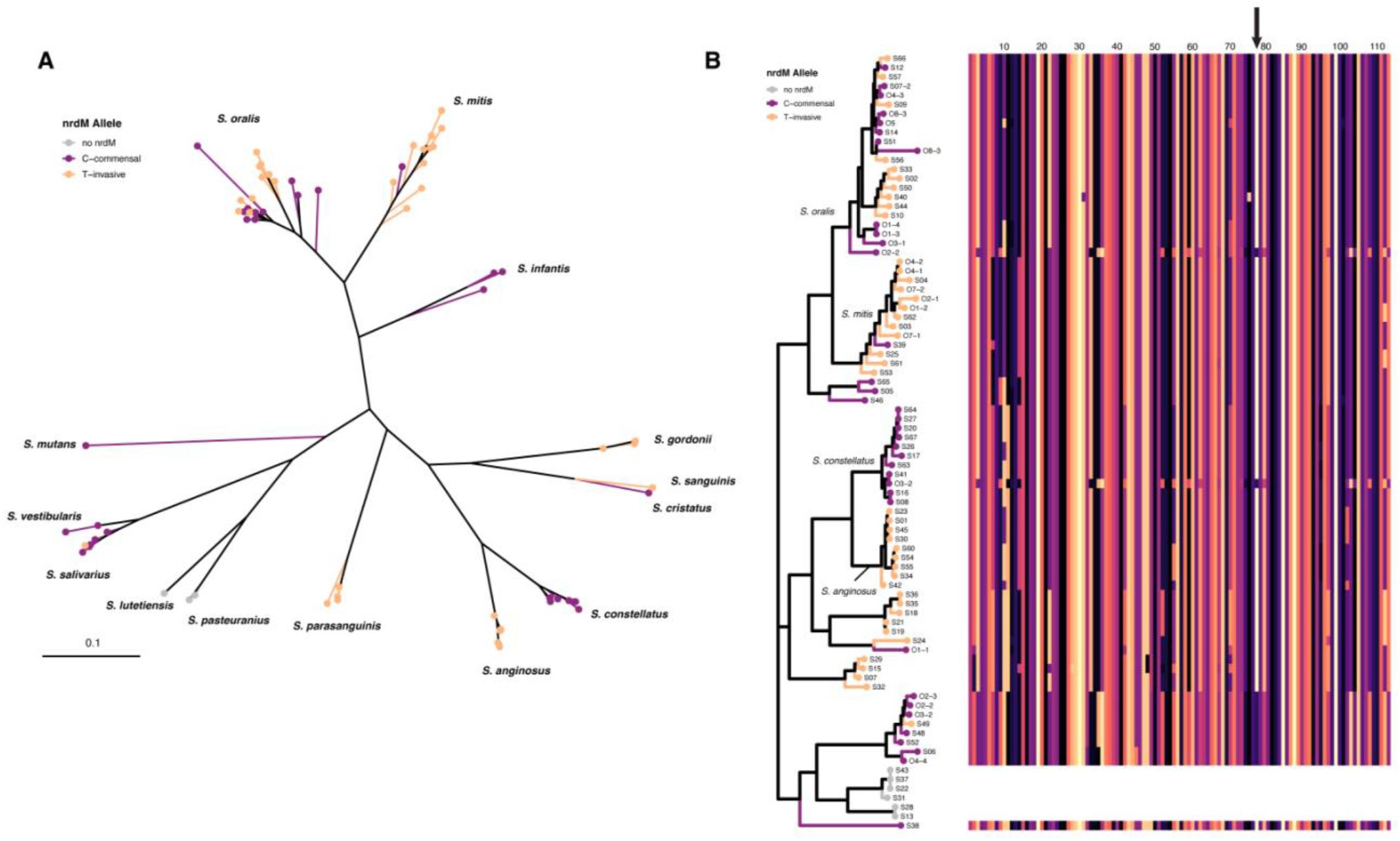
**A)** Homologous *nrdM* genes identified in all newly sequenced VGS isolates with alleles from our variant of interest plotted on a phylogeny made from GyrB sequences. Only 2 species in our sample lacked *nrdM (S. lutetiensis* and *S. pasteuranius*) both of which are in the bovis group and are shown in grey. All other species contained *nrdM* and were conserved at the position of interest. **B)** Multiple sequence alignment (MSA) of NrdM plotted next to the *gyrB* phylogeny (same tree as in A) where tips have been colored by *nrdM* allele. Scale on top of the MSA represents length in amino acids, the position of the synonymous mutation of interest (I78I) indicated with an arrow. Clade labels shown for the most common species: *S. oralis, S. mitis, S. anginosus* and *S. constellatus*.

### Deletion of *nrdM* does not affect *in vitro* growth

To investigate the impact the *nrdM* orphan allele has on growth in the two infective endocarditis strains *S. oralis* 1648 (SO48) and mitis group *Streptococcus* 1643 (SM43), we deleted *nrdM*, locus tags MP387_03665 and FD735_06230, respectively, from each strain, via homologous recombination (Table 1). SO48 possesses the *nrdM-C* (“C” = commensal) allele, and SM43 possesses the *nrdM*-I (“I” = invasive) allele. Knockout mutants were confirmed via Sanger sequencing. No difference in growth was observed for knockout mutants compared to wild-type in Todd-Hewitt broth (THB) (Figure S6). Further, *nrdM* allele swaps from SO48 and SM43 were placed back into the genome of SO48Δ*nrdM* and SM43Δ*nrdM* to generate Δ*nrdM*::*nrdM*-I or Δ*nrdM*::*nrdM*-C strains in each strain background (Table 1). Since the presence of human serum has been shown to impact mitis group streptococci physiology (*45*, *46*), we assessed the impact of *nrdM* during growth with human serum by performing growth curves of allele replacement strains in chemically defined medium supplemented with 5% v/v human serum (Figure S6). No difference in growth profile was observed between alleles in either strain background. Thus, the *nrdM* locus does not impact growth in rich laboratory medium or chemically defined medium supplemented with 5% v/v human serum *in vitro*.

**Table 1:** Strains used in this study.

| Organism | Strain | Description | Ref |
| --- | --- | --- | --- |
| Mitis group streptococci | 1643 (SM43) | Wild-type infective endocarditis isolate | (19) |
|  | SM43ΔNrdM | SM43 with clean deletion of *nrdM*, FD735_06230 | This work |
|  | SM43:NrdM-I | SM43 with SM43 *nrdM* allelic replacement into SM43Δ*nrdM* | This work |
|  | SM43:NrdM-C | SM43 with SO48 *nrdM* allelic replacement into SM43Δ*nrdM* | This work |
| *S. oralis* | 1648 (SO48) | Wild-type infective endocarditis isolate | (19) |
|  | SO48ΔNrdM | SO48 with clean deletion of *nrdM*, MP387_03665 | This work |
|  | SO48:NrdM-I | SM43 *nrdM* allelic replacement into SO48Δ*nrdM* | This work |
|  | SO48:NrdM-C | SO48 *nrdM* allelic replacement into SO48Δ*nrdM* | This work |

### *nrdM* has no impact on growth in a simulated infective endocarditis vegetation model

Finally, the allele swapped strains were investigated using the pharmacological *in vitro* simulated infective endocarditis vegetation model (SIEVM). We reasoned that if *nrdM*-I confers enhanced fitness in simulated endocardial vegetations, we would observe significantly different vegetation CFUs for SO48Δ*nrdM*::*nrdM*-I versus SO48Δ*nrdM*::*nrdM*-C. The SM43 strain was similarly tested, to assess the effect of *nrdM* allele swapping in a closely related but different (i.e. non-*oralis*) genetic background. Strains were inoculated into vegetation clots at ~ 1 x10^8^ CFU, incubated in the chemostat model with Mueller Hinton Broth (MHB) in parallel for 48hrs (see Material and Methods). At 4, 24, and 48 h post inoculation, 4 clots from each model were removed for CFU enumeration (Figure 6). No significant difference was observed during growth in the SIEVM between *nrdM* allele swap in either strain background. Ultimately, these data and the growth curve data together show that under these *in vitro* conditions, *nrdM* is not essential, and no phenotype could be assigned to either of the *nrdM* alleles.

**Figure 6:**
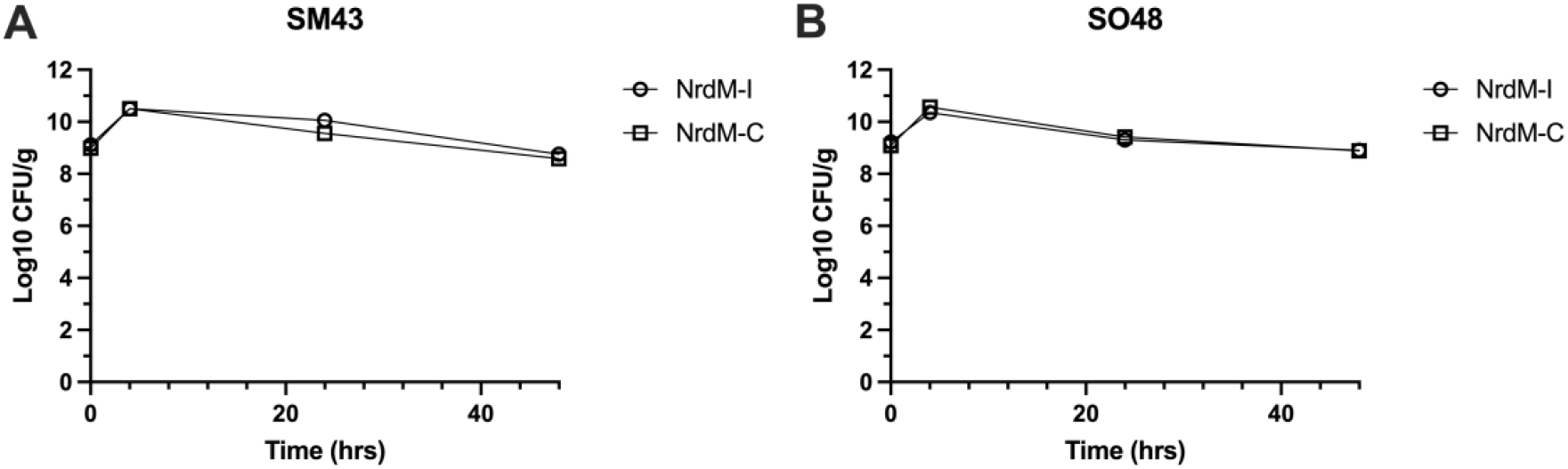
Simulated infective endocarditis vegetation model of *nrdM* allele swapped strains. Survival of either **A)** SM43 or **B)** SO48 harboring either the infectious *nrdM*-I allele or the commensal *nrdM*-C allele during growth in vegetations. Biological triplicates were performed for each strain with four technical replicates per time point. Mean and SEM indicated. No significant difference was observed under these conditions. Two-way ANOVA with Sidak’s multiple comparisons test.

## Discussion

The viridans group streptococci (VGS) are a large collection of closely related *Streptococcus* spp. that inhabit the oral cavity and gastrointestinal and genitourinary tracts of humans as commensals, but can invade other tissues to cause severe diseases such as bacteremia and infective endocarditis (*1*). Our understanding of the pathogenesis of VGS disease is limited by lack of knowledge surrounding the genetic and environmental conditions that facilitate a switch from commensalism to pathogen. Additionally, clinical detection and identification of bacterial infections is critical for patient care and recovery, especially in neutropenic and immunocompromised patients. A major etiological agent of disease in this patient demographic are the VGS, especially *S. mitis* and *S. oralis*. Yet, clinical methods of speciation are still largely inaccurate, and even newer methods like MALDI-TOF MS struggle to differentiate between closely related species such as *S. oralis* and *S. mitis* (*7*). In this study, we collected and whole genome-sequenced a variety of invasive and non-invasive VGS isolates, compared bioinformatic methods for delineating closely related species, and identified a SNP that may predispose certain isolates of *S. oralis* to invasive disease.

### Complex population structure of *S. oralis*

We characterized the population structure of 81 *S. oralis* isolated from healthy oral microbiota and from blood, and found high levels of diversity, both in the core (Figure 2B-C) and accessory genomes (Figure 2A,D). We found that using our methods, previously described sub-species of *S. oralis* corresponded to sub-clades we identified in a phylogeny inferred from core genome sequences (Table S2). We also identified *subsp. oralis* as being the least diverse of the three subspecies, with respect to both gene content and sequence variation in the core genome (Figure 2). We assert that recombination is likely the primary mechanism for generating and maintaining diversity in this species, as 99.9% of the *S. oralis* core genome has been affected by recombination (Figure 3). The rarity of a given recombinant fragment in our sample also indicates that *S. oralis* isolates are likely participating in HGT with diverse species – made possible by the fact that they inhabit complex communities. We observed a mean piN/piS value across *S. oralis* core genes of 0.039 which is an order of magnitude lower than comparator species such as *S. aureus* for which values of 0.32 (*47*) and 0.55 (*48*) have been reported, and lung-colonizing *P. aeruginosa* from cystic-fibrosis patients that have a mean of 0.14. (*49*). This indicates that *S. oralis* core genes are under strong purifying selection. We recently identified a similar phenomenon among environmental isolates of *Mycobacterium abscessus* that are also highly recombinogenic and manifest high amounts of synonymous diversity in their core genomes (*50*). This suggests that like *M. abscessus, S. oralis* inhabits environments alongside diverse microbial species where high rates of recombination enable ready acquisition of novel genetic material and rapid removal of deleterious mutations.

### Switching from commensal to pathogen

Our identification of extremely high recombination rates within *S. oralis* points to the unique genomic features characterizing this VGS species. High rates of recombination allow the species to adapt to fluctuating environments encountered within the human body, and potentially enable invasion of new niches, including pathogenic niches. We used three different GWAS methods that replicably identified a strong association between a synonymous SNP within an undescribed protein, NrdM, and invasive disease (Figure 4, Figure S2). The association was robust across diverse isolates of all three subspecies (Figure S3). We also found that *nrdM* was conserved among *S. oralis* isolates (Figure S4) and is one of only ~440 genes conserved across all VGS species in our sample indicating that it is likely to play an important, and perhaps a similar, role across species. Utilizing *in vitro* growth in the presence of human serum and a simulated infective endocarditis vegetation model, we compared *nrdM* knockout strains and allele-swapped invasive and commensal isolate alleles in two strain backgrounds. No phenotype was observed under the tested conditions, suggesting that if NrdM is contributing to invasive disease, it requires different conditions (possibly specific to the *in vivo* environment) for a potential phenotype to be observed. NrdD, which shares the same ATPase cone domain as NrdM, is required for anaerobic growth in *S. sanguinis* (*39*), and, notably, disruption of this gene results in attenuated virulence (*51*). Long thought to have neutral fitness effects, synonymous mutations are increasingly recognized as having significant effects on bacterial fitness, for example by impacting gene expression and protein folding (*52–56*). We believe that further study of this synonymous variant could reveal fitness effects and that adaptation within *nrdM* may affect a multitude of phenotypes that aid in transition from the oral cavity to the bloodstream, although further investigation is necessary.

A major hurdle to successful genomic analyses in VGS is the correct classification of the different species. It is possible that incorrect species identification has contributed to the lack of clear results from studies looking at the genetic determinants of virulence in VGS species. One such study by Rasmussen *et. al*. used a sample of both *S. mitis* and *S. oralis* to search for known virulence factors, which they found at varying frequencies, indicating that genetic differences may be responsible for variability in virulence among strains (*57*). Similar to our sample of *S. oralis* isolates (Figure 2B), studies of the closely related VGS *S. sanguinis* and *S. gordonii* were unable to identify phylogenetic patterns based on invasive disease (*58*, *59*). These studies, however, were unable to identify specific genomic variants associated with invasiveness. What our work and the work of others indicates, is that virulence properties differ between VGS species, despite being so closely related and often causing similar disease. This is further illustrated by the fact that although *nrdM* is conserved in *S. mitis*, the association between the invasive allele identified in *S. oralis* and pathogenicity in *S. mitis* is not significant (data not shown). Going forward, analyses using larger samples of well-defined, individual species will provide better resolution for identifying the genetic determinants of virulence in the VGS.

## Materials and methods

### Collection of clinical Isolates

Clinical strains obtained from routine blood cultures and identified as viridans group streptococci were stored in the microbiology laboratory at each hospital (Methodist Health System [MHS] and University of Mississippi Medical Center [UMMC]). Respective site investigators reviewed patient clinical information to confirm invasive infection (i.e., bacteremia or endocarditis), then de-identified specimens and provided blinded isolates to the University of Texas at Dallas (UTD) laboratory for study. Specimens were not included for further analysis if the isolate was deemed a contaminant and did not require antibiotic therapy as determined by the treating physician. Site investigators obtained approval from respective Institutional Review Boards (MHS UTD IRB 18-121 and UMMC 2018-0068).

### Processing of clinical isolates

Clinical isolates were struck on Mitis-Salivarius Agar (MSA) (BD Bacto and incubated overnight in 37°C and 5% CO_2_. MSA plates were observed for homogenous colony morphology, and a single colony was inoculated into 10 mL THB for overnight growth at 37°C and 5% CO_2_. If more than one colony morphology was identified on MSA plates, broth cultures were made from each colony morphology. Overnight cultures were stored at −80°C in 25% glycerol. The remaining culture volume was pelleted at 4,280 x *g* in a Sorvall RC6+ floor centrifuge and genomic DNA was extracted using Qiagen DNeasy Blood and Tissue Kit per manufacturer protocols, with minor modifications as described in (*6*).

### Collection of oral swab samples

Oral swabs were obtained from healthy adult volunteers at the UTD campus (UTD IRB 17-170). The following exclusion criteria were applied: previous history of bacteremia or endocarditis; recent antibiotic exposure (prior 30 days); history of periodontal disease; personal or family history of immunocompromise. No participants were excluded based on these criteria. The volunteer was asked to rinse their mouth with sterile saline, and then self-swab their teeth and tongue with a sterile swab (Puritan). Swabs were stored at 4°C until processing.

### Processing of oral swabs

Oral swabs were processed as described above. Briefly, swabs were struck onto MSA plates and grown overnight. After incubation, the MSA plates were observed and colony morphologies consistent with *Streptococcus mitis* and *S. oralis* were selected for overnight growth in THB. In addition, approximately 3 random colonies of different morphologies were also selected for overnight growth. Cultures were processed as described above.

### 16S rRNA sequence analysis and GyrB typing

PCR reactions were performed using Taq polymerase (New England Biolabs) with primer sequences in Table S3. 16S rRNA genes were amplified using universal primers 8F and 1492R (*60*). The DNA Gyrase B gene (*gyrB*) was amplified using previously reported primers (Table S3). PCR reactions were analyzed by agarose gel electrophoresis and purified using the GeneJET PCR purification kit (Thermo Fisher) per the manufacturer protocols. Products were sequenced at the Massachusetts General Hospital DNA Core. 16S rRNA sequences were trimmed using Geneious R11 (https://www.geneious.com) allowing a maximum of 10 low quality bases and 6 ambiguities. Trimmed sequences were used as queries for NCBI Blastn against the 16S ribosomal RNA sequences (Bacteria and Archaea) database and species were assigned only when the forward and reverse sequencing reactions had the same top Blastn result (Table S1). GyrB nucleotide sequences were translated, and amino acid sequences pairwise aligned to *Streptococcus mitis* ATCC 49456 GyrB (locus tag SM12261_0755). Amino acid variations were identified using the method of Galloway-Peña et al. (*11*).

### Ilumina sequencing

Sequencing was performed at The University of Texas at Dallas Genome Core using Illumina Nextseq 500 platform with a mid-output 300 cycle of 2 × 75 bp paired-end reads for clinical isolates S1-S31, or 2 × 150 bp paired-end reads for all other isolates.

### Genome assembly & annotation

Using the raw sequencing data from newly sequenced clinical and oral isolates (Table S1), species identification was additionally performed using Kraken2 (*20*) Raw data was quality checked and trimmed using FastQC v0.11.8 (*61*) and TrimGalore v0.6.4, (http://www.bioinformatics.babraham.ac.uk/projects/trim_galore), respectively. Contigs were assembled using SPAdes v3.13.0 with default parameters (*62*). Assemblies were checked for quality using Quast v5.0.2 (*63*) filtering out contigs shorter than 500bp or with coverage lower than 5x, as well as confirming all assemblies had a N50 > 50,000bp. Contigs were annotated using Prokka v1.13.3 (*64*) before a pangenome analysis was performed with Roary v3.12.0 using a blastp identity threshold of 75% (*31*). Using a nucleotide sequence alignment of GyrB (as clustered by Roary) a phylogenetic tree was made using FastTree v2.1.9 (*65*) and visualized in R with ggtree (*66*). GyrB typing was confirmed using these sequences with a custom script (code available at https://github.com/myoungblom/VGS_GWAS.git) assigning species based on the scheme outlined by Galloway-Peña et al. (*11*). *nrdM* sequences from all newly sequenced VGS isolates were identified with Blastp using the *S. oralis* NrdM amino acid sequence as the query sequence.

### *S. oralis* genome collection for GWAS analyses

From our sample of clinical and oral isolates, *S. oralis* made up the largest part of our sample so we decided to proceed with analyses of just this species. To create a dataset large enough for a powerful genome-wide association study (GWAS) we identified all *S. oralis* and *S. mitis* isolates from NCBI (NCBI SRA & Assembly databases accessed June 2019) with the proper metadata indicating they were isolated from the mouth (e.g oral cavity, dental plaque, dental biofilm, etc) or from blood (e.g. infective endocarditis, bloodstream infection, blood, etc). We assumed isolates uploaded to NCBI with various “oral” sources were all commensal and all those from “blood” were from an invasive infection (Table S2). We chose to start with both *S. oralis* and *S. mitis* because these species are so closely related, they are often mistaken for each other and uploaded to NCBI under the wrong species (Table S2) as has previously been reported (*10*). We pulled out the true *S. oralis* isolates using Kraken and GyrB typing as described above. Samples for which raw sequence data was available were assembled as described above, and then annotated with the remainder of assemblies downloaded from NCBI (Table S2). Note that since the time of data collection, three of the assemblies used in this dataset have been suppressed (Table S2). We then performed a pangenome analysis on the *S. oralis* sample as described above (using a Blastp identity threshold of 95% and Prank to align core genes) and a phylogenetic tree was inferred from the resulting core genome alignment using RAxML v8.2.3 (*67*) and visualized in R with ggtree (*66*).

### Recombination analyses

We identified recombinant fragments in the *S. oralis* core genome using Gubbins v2.4.1 with default parameters (*35*). Recombinant fragments were visualized alongside the core genome phylogeny using Phandango (*68*). We used ClonalFrameML v1.11 (*36*) with default parameters to estimate *r/m*.

### Population genetics statistics

Pairwise average nucleotide identity (ANI) values of *S. oralis* core genome sequences were calculated with OrthoANI (*69*). piN/piS values were calculated for all pairwise combinations for each *S. oralis* core gene were calculated using Egglib (*70*), then the average piN/piS value for each gene was calculated.

### Rarefaction & accumulation plots

Rarefaction and accumulation curves for the *S. oralis* subspecies were calculated from Roary gene presence absence files. Briefly, separate pangenome analyses were performed as described above for each individual subspecies and then each dataset was iteratively subsampled to the size of the smallest dataset (n=25) and the median number of core and total genes was plotted from all iterations.

### GWAS

We first queried for genetic associations with the ‘invasive’ phenotype in our dataset by identifying accessory gene content significantly associated with invasiveness using Scoary v1.16.6 (*38*). We then performed a preliminary GWAS of core genome variants using an F_ST_ outlier analysis. Briefly, a VCF file containing all core genome variants was made using SnpSites v2.0.3 (*71*) and reformatted using a custom script (code available https://github.com/myoungblom/VGS_GWAS.git). Then we calculated Weir and Cockerham’s F_ST_ for bi-allelic SNPs using vcflib (https://github.com/vcflib/vcflib). Using a custom script (code available https://github.com/myoungblom/VGS_GWAS.git) we permuted the phenotypes in this analysis 100x and used the maximum F_ST_ value observed in the null distribution as a cutoff to identify significant F_ST_ outliers. To validate the results of our F_ST_ outlier analysis we also used two GWAS programs designed specifically for use with microbial genomes: treeWAS (*40*) that corrects for the presence of recombination, and BugWAS (*41*) that identifies lineage effects and controls for population structure. TreeWAS was run using the recombination-adjusted phylogenetic tree made with Gubbins (see above) using 10x the number of SNPs in the core genome for the parameter “n.snps.sim”. BugWAS was run using default parameters.

### Homoplasy analysis

Homoplasy analysis was performed using TreeTime v0.9.0-b.2 (*44*) with default parameters.

### Deletion of *nrdM*

Knockout constructs of *nrdM* in SM43 (Locus ID FD735_06230) and SO48 (Locus ID MP387_03665) were generated as previously described (*45*). Briefly, linear constructs were generated by amplifying ~2 kb regions upstream and downstream of *nrdM* using Phusion polymerase (Thermo Fisher) using primers in Table S3. SOEing PCR was used to stitch fragments together and the amplified product was assessed via agarose gel electrophoresis. Gel extraction was performed using the QIAQuick Gel Extraction kit (Qiagen). Linear constructs were transformed by natural transformation as described in (*45*). Transformation plates were incubated overnight, and putative transformant colonies were screened via PCR for the *nrdM* deletion.

### *nrdM* allele swaps

Allele swap strains were generated using the same strategy as the deletion, except the linear construct contained either the *nrdM*-I or *nrdM*-C allele coupled with the flanking regions for the respective strain. The linear construct was transformed into SM43Δ*nrdM* and SO48Δ*nrdM*. The allele swap region in transformants was amplified using primers in Table S3. Products were sequenced for validation of the allele swap (Massachusetts General Hospital DNA Core).

### Growth curves

Growth curves in THB were performed in biological duplicate. Wild-type and *ΔnrdM* strains were grown overnight as described, then diluted to OD_600nm_ 0.05 in approximately 12 mL THB. OD_600nm_ was monitored every hour using a Thermo Scientific Genesys 30 spectrophotometer. For growth curves in the presence of human serum, biological triplicate overnight cultures were grown in streptococcal defined medium (*45*, *46*, *72*) and diluted to OD_600nm_ 0.1 in defined medium supplemented with 5% v/v human serum (Sigma-Aldrich). OD_600nm_ was monitored at 3, 6 and 24 hrs as described above.

### Simulated Infective Endocarditis Vegetation Model (SIEVM)

Strains were inoculated from freezer stocks into 5 mL Mueller Hinton Broth (MHB) (BD Bacto) and incubated overnight at 37°C and 5% CO_2_. 1 mL was expanded into 100 mL pre-warmed MHB and incubated overnight as described above. The SIEVM was set up and performed as previously described (*19*) using human blood products from the American Red Cross (UTD IRB 19MR0160). Briefly, 500 μL of pooled human cryoprecipitate (American Red Cross), 50 μL ~2 TIU/mL aprotinin (Sigma-Aldrich), ~100,000 human platelets (American Red Cross), and 10^8^ CFU/g bacteria were combined in a sterile Eppendorf tube and vortexed. Sterile monofilament line was positioned before 100 μL of ~2 KU/mL high activity bovine thrombin (Sigma-Aldrich) was added to congeal the vegetation. Vegetations were placed into the glass apparatus in a 37°C water bath, and MHB was pumped through the model at a pre-calibrated rate of 0.4 mL/min. Four vegetations were removed at designated time points for each strain, weighed, removed from the monofilament line and placed in 1.25% Trypsin solution (Sigma-Aldrich) in sterile screw cap microcentrifuge tubes (Fisher Scientific) with ~5-8 2.7 mm glass beads (BioSpec). Clots were homogenized for 10-15 min horizontally on a vortex before serial dilution and plating on THB agar plates for enumeration. CFU/g was calculated by multiplying the observed CFU/mL by the net weight of the vegetation. SIEVM was performed in biological triplicate for each strain, with four vegetations per time point per strain.

### *S. oralis* 1648 hybrid genome assembly

Pacific Biosciences single molecule real time (SMRT) sequencing was performed by the Johns Hopkins Genome Core. The SO48 whole genome was assembled using the Unicyler assembly pipeline (*73*) combining SMRT long reads generated in this study and Illumina reads previously generated for SO48 (accession PRJNA354070; (*74*)).

### Accession numbers

The SO48 whole genome sequence generated in this study has been deposited in Genbank under the accession number CP094226. Genome constructs and Illumina and SMRT sequence reads generated in this study have been deposited in the Sequence Read Archive under the BioProject accession PRJNA817585, see Table S1.

## Acknowledgements

This work was supported by R21 AI130666 from the National Institutes of Health and the Cecil H. and Ida Green Chair in Systems Biology Science to K.L.P., and by the National Science Foundation Graduate Research Fellowship to M.A.Y under grant number DGE-1747503. The content of this work is solely the responsibility of the authors and does not necessarily represent the official views of the National Institutes of Health.

