## Supplemental figures for "Comparative genomics of *Streptococcus oralis* identifies large scale homologous recombination and a genetic variant associated with infection"

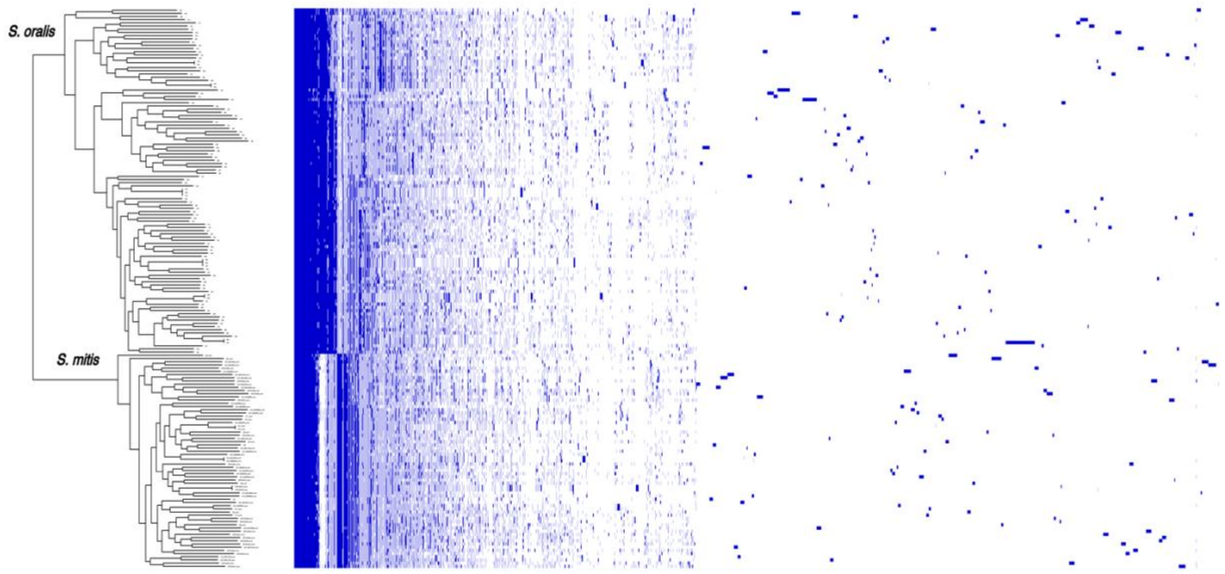


**Figure S1:** Core genome phylogeny of all *S. oralis* and *S. mitis* isolates (from newly sequenced samples and from NCBI) alongside a matrix of gene content. Clear delineation between *S. oralis* and *S. mitis* can be seen both in the core genome sequence as well as in core and accessory gene content differences between the species. Pangenome analysis performed using Roary and visualization with Phandango.

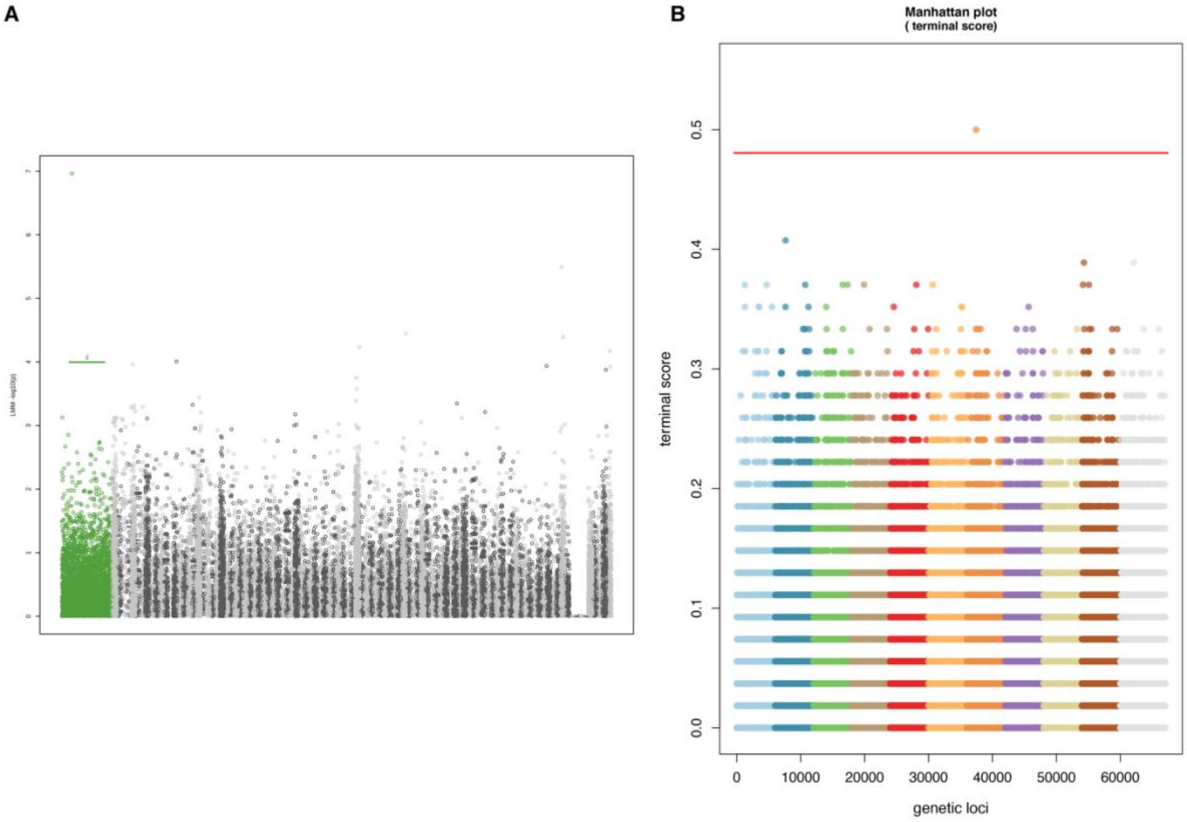


**Figure S2:** GWAS results confirming results of $F_{ST}$ outlier analysis. BugWAS (left) and TreeWAS (right) results each showing a single significant variant associated with invasiveness: the same variant in *nrdM* identified in the $F_{ST}$ outlier analysis.

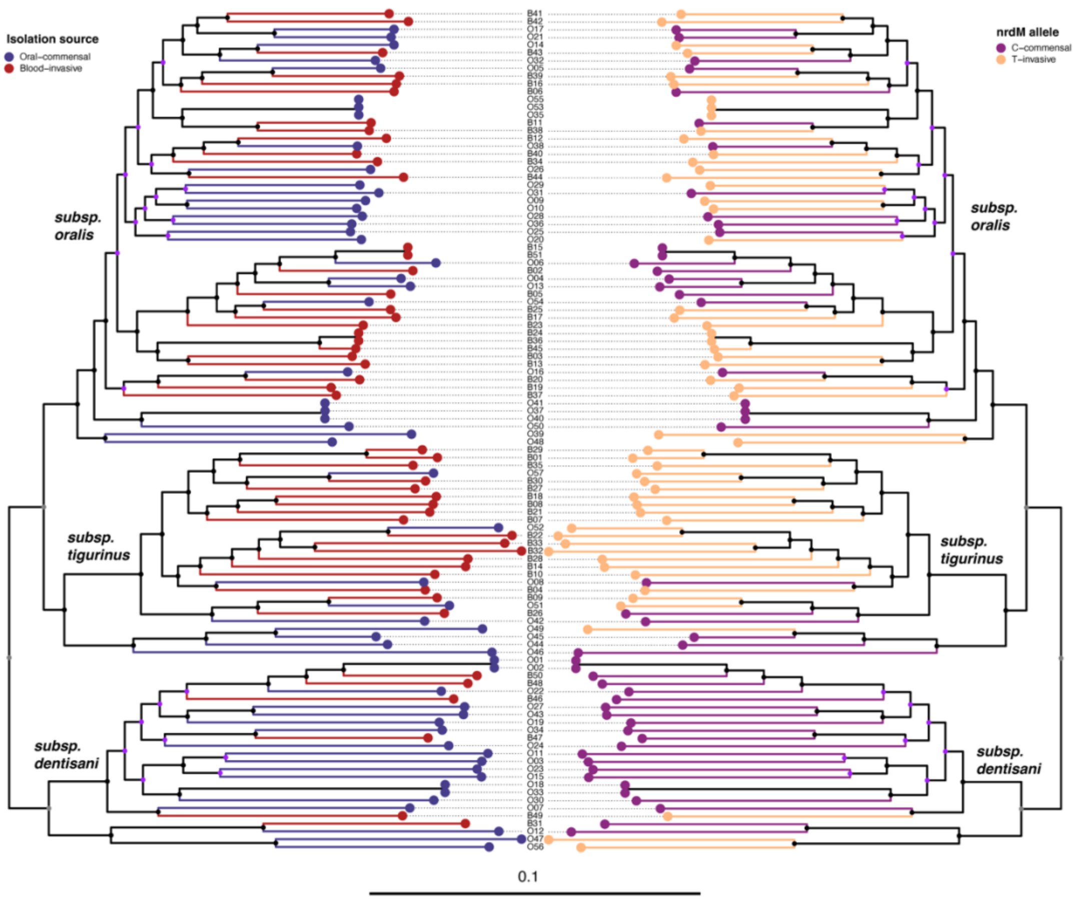


**Figure S3:** Comparison between isolation source (oral commensal or blood/invasive infection) and *nrdM* allele (C or T) shown on mirrored core genome phylogenies. Tree tips connected by dotted lines have convergent genotype-phenotype combinations (*i.e.* the *nrdM* allele associated with invasiveness in an isolate from an invasive infection).

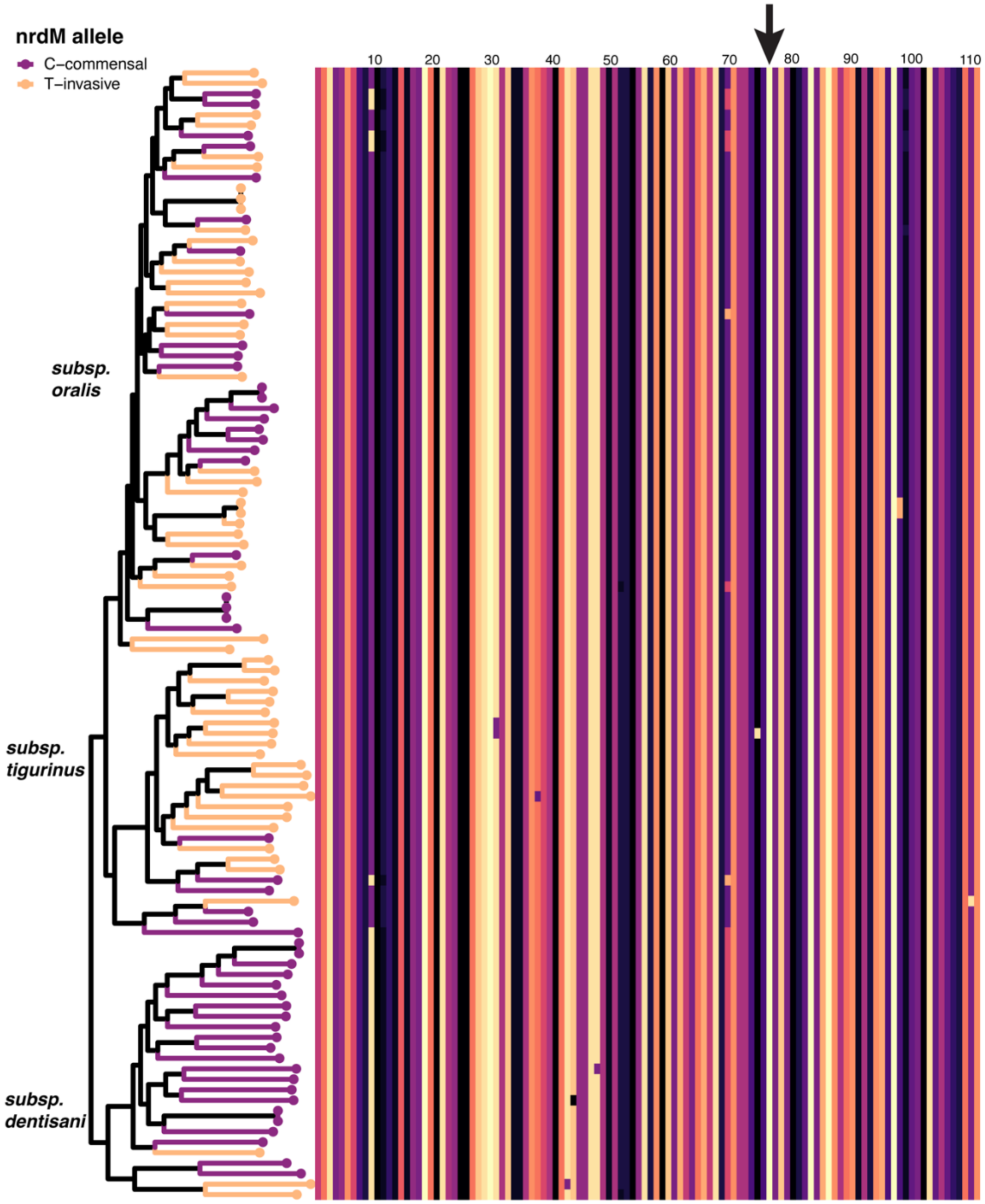


**Figure S4:** Multiple sequence alignment (MSA) of NrdM plotted next to the *S. oralis* core genome phylogeny where tips have been colored by *nrdM* allele. Scale on top of the MSA represents length in amino acids, the position of the synonymous mutation of interest (I78I) indicated with an arrow.

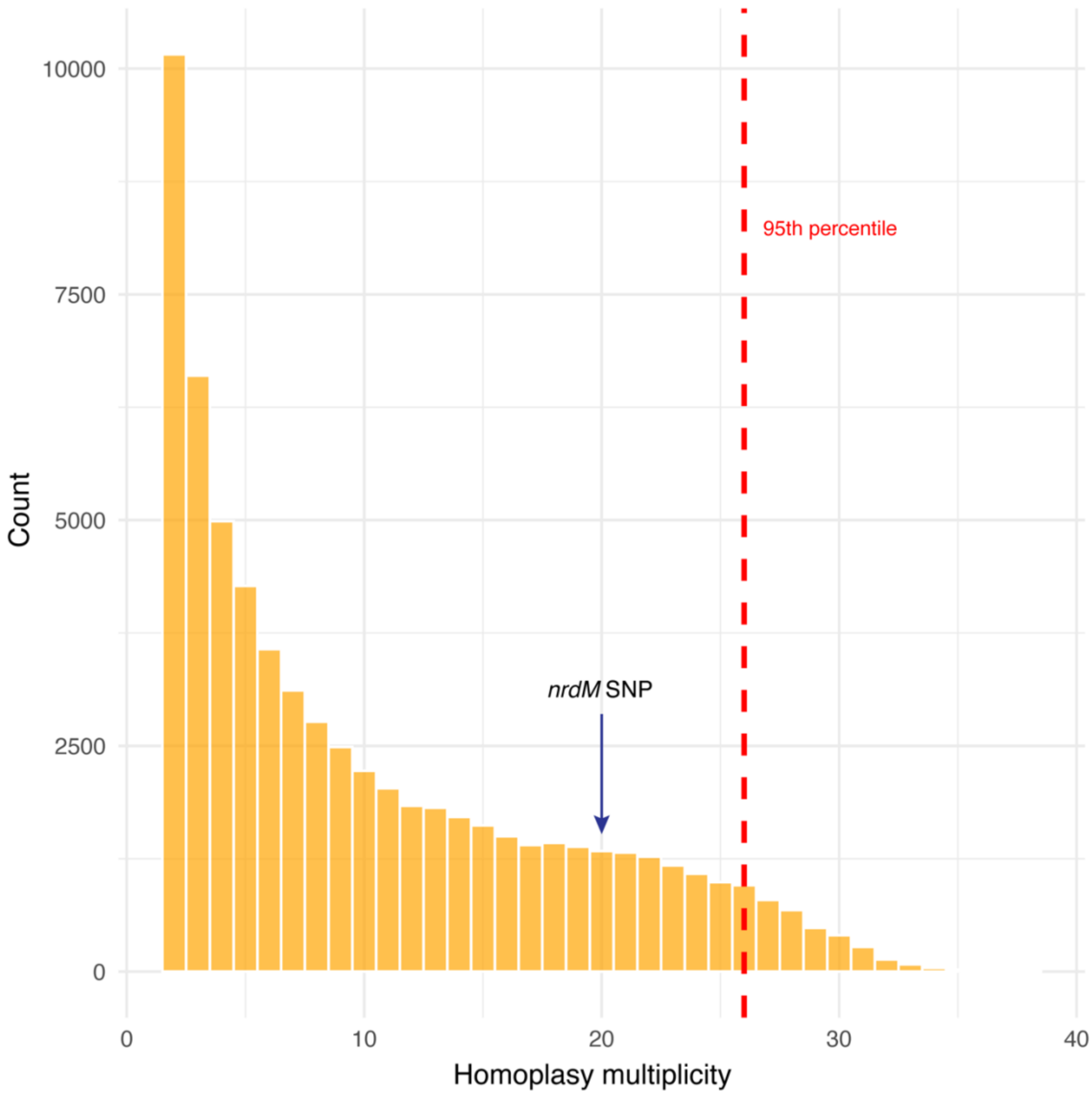


**Figure S5:** Histogram of mutation multiplicity for all homoplastic mutations on the *S. oralis* core genome phylogeny. Red dotted line shows 95$^{th}$ percentile cutoff and the multiplicity of the *nrdM* SNP (20) indicated by the black arrow.

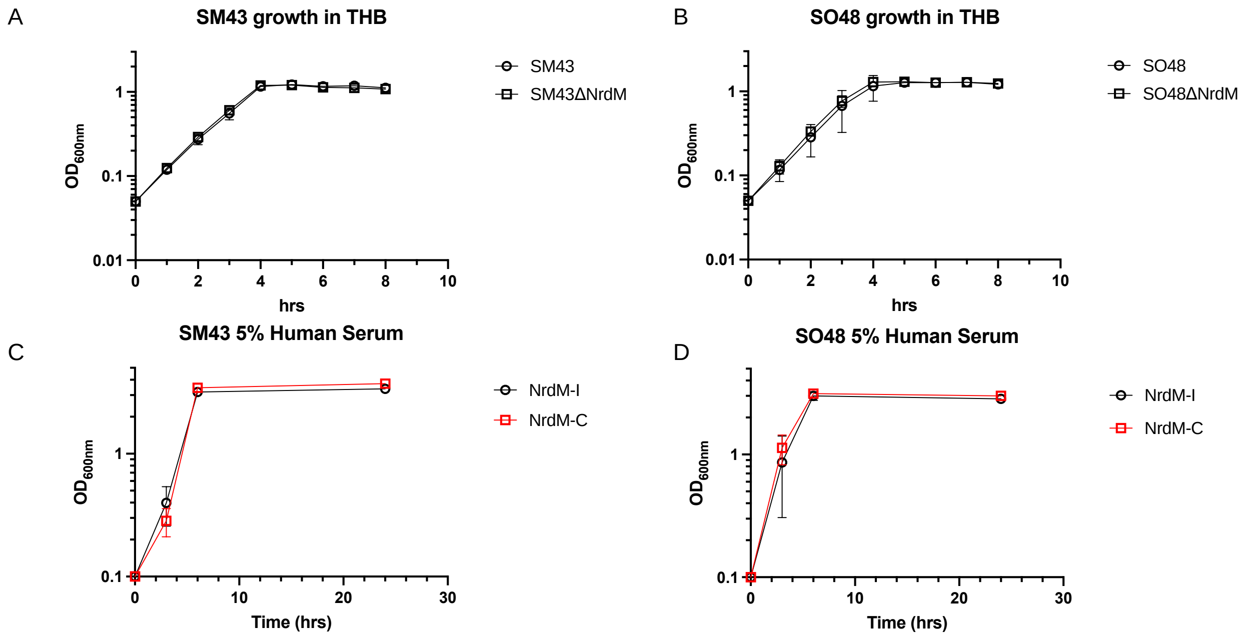


**Figure S6: Growth curves of mutant strains**. Growth curves of wildtype and Δ*nrdM* strains of A) SM43 and B) SO48 performed in Todd-Hewitt Broth, in biological duplicate. OD600 readings were performed every hour for 8 hrs. Mean and SD indicate. Growth of NrdM-I and NrdM-C alleles in C) SM43 and D) SO48 strain backgrounds when grown in streptococcal defined medium supplemented 5% v/v human serum. Serum growth was performed in biological triplicate and OD600 readings were taken at 0, 3, 6, and 24 hrs. Mean and SEM indicated.

**Table S1:** Newly sequenced VGS isolates

**Table S2:** *S. oralis* GWAS Strains

**Table S3**: Primers used in this study
