## Supplementary material for "Comparative genomics of *Streptococcus oralis* identifies large scale homologous recombination and a genetic variant associated with infection": Table S3

Table S3. Primers used in this study

| Primer | Sequence (5'-3') | Use |
|---|---|---|
| 43_A1 F | TTGTGAAAATCCCTAGAACG | SM43 upstream of *nrdM* region amplification |
| 43_K1 F | AACCTTATGCAACGTTAAGAAAAAGAGTGG | SM43 downstream of *nrdM* region amplification for clean deletion |
| 43_D1 O F | GAAAACTATCAACTGACAGC | SM43 upstream of *nrdM* for screening |
| 43_D2 F | ATCCAGATAAAATTTACCAAGC | Internal primer of SM43 *nrdM* |
| 43_I1 F | CGACATTGATCTAACCAAAACC | SM43 upstream of *nrdM* region outside recombination arms |
| 43_K1 R | TTTTCTTAACGTTGCATAAGGTTCTCC | SM43 upstream of *nrdM* region amplification for clean deletion |
| 43_A1 R | TTGTTACCTTAGTTAAAAGAATAGG | SM43 downstream of *nrdM* region amplification |
| 43_D2 O R | CCTCAACTGTACAAAAAGC | SM43 downstream of *nrdM* for screening |
| 43_D1 R | GATAAGAGATATAGTGTTCTGC | Internal primer of SM43 *nrdM* |
| 43_I1 R | CAAGGCAGACTTTGATGC | SM43 downstream of *nrdM* region outside recombination arms |
| 48_A F | GCAAGAATTTTTGTGACAAGG | SO48 upstream of *nrdM* region amplification |
| 48_K F | CCTTATGCAACGTTAAGAAAAAGAGTGG | SO48 downstream of *nrdM* region amplification for clean deletion |
| 48_D1 O F | GAAAACTATCAACTGACAGC | SO48 upstream of *nrdM* for screening |
| 48_D2 F | AAACGTAATGGAGAAATTGC | Internal primer of SO48 *nrdM* |
| 48_I F | TAAAATCAATAAAGAGAGCTACGG | SO48 upstream of *nrdM* region outside recombination arms |
| 48_K R | TTTTCTTAACGTTGCATAAGGTTCTCC | SO48 upstream of *nrdM* region amplification for clean deletion |
| 48_A R | GGTTGACTTTTCTTTCTGAATTAG | SO48 downstream of *nrdM* region amplification |
| 48_D2 O R | CCTCAACTGTACAAAAAGC | SO48 downstream of *nrdM* for screening |
| 48_I F | TAAAATCAATAAAGAGAGCTACGG | Internal primer of SO48 *nrdM* |
| 48_I R | AGCTATCCACCCAACCG | SO48 downstream of *nrdM* region outside recombination arms |
| 1492R | CGGCTACCTTGTTACGACTT | 16S rRNA universal primer |
| 8F | AGAGTTTGATCCTGGCTCAG | 16S rRNA universal primer |

| | | |
|---|---|---|
| UpRep_R | ATTACTTGCATAAGGTTCTCCTTTATTCTTTG | Amplifies upstream fragment of *nrdM* |
| Rep_F | AGGAGAACCTTATGCAAGTAATCAAACG | Amplifies *nrdM* gene |
| Rep_R | CACTCTTTTTCTTAACGGATTTGTTCAAAATG | Amplifies *nrdM* gene |
| DwnRep_F | AATCCGTTAAGAAAAAGAGTGGGAT | Amplifies downstream fragment of *nrdM* |
| GyrB_F | CAAGGTTTCCGTACAGC | Amplifies *gyrB* |
| GyrB_R | GCTTCTTGGAGTTTAATTTCTTG | Amplifies *gyrB* |
